# Maternal immune activation causes age-specific changes in cytokine receptor expression in offspring throughout development

**DOI:** 10.1101/490466

**Authors:** Myka L. Estes, Bradford M. Elmer, Cameron C. Carter, A. Kimberley McAllister

## Abstract

Maternal infection is a shared environmental risk factor for a spectrum of neuropsychiatric disorders and animal models of maternal immune activation (MIA) exhibit a range of neuropathologies and behaviors with relevance to these disorders. In particular, MIA offspring show chronic, age- and region-dependent changes in brain cytokines, a feature seen in postmortem studies of individuals with neuropsychiatric disorders. These MIA-induced alterations in brain cytokines may index biological processes underlying progression to diagnosable neuropsychiatric disorders. However, cytokines signal through specific cytokine receptors to alter cellular processes and it is the levels of those receptors that are critical for signaling. Yet, it remains unknown whether MIA alters the expression of cytokine receptors in the brains of offspring throughout postnatal development. Here, we measured the expression of 23 cytokine receptors in the frontal cortex of MIA and control offspring from birth to adulthood using qPCR. MIA offspring show dynamic oscillating alterations in cytokine receptors during sensitive periods of neural growth and synaptogenesis. Of the many cytokine receptors altered in the FC of MIA offspring, five were significantly changed at multiple ages at levels over 2-fold relative to controls—*Il1r1, Ifngr1, Il10ra, Cx3cr1 and Gmcsfr*—suggesting persistent dysfunction within those pathways. In addition to facilitating immune responses, these cytokine receptors play critical roles in neuronal migration and maturation, synapse formation and elimination, and microglial function. Together with previously reported changes in cytokine levels in the brains of MIA offspring, our results show a decrease in cytokine signaling during the peak period of synaptogenesis and spine formation and an increase during periods of activity-dependent development and early adulthood. Overall, the oscillating, age-dependent cytokine receptor alterations in the FC of MIA offspring identified here may have diagnostic and therapeutic value for neuropsychiatric disorders with a neuro-immune etiology.

**Research Highlight:** Maternal immune activation leads to long-lasting, oscillating changes in cytokine receptors in the frontal cortex of offspring throughout development.

## Introduction

Psychiatric disorders are the leading cause of disability worldwide and exact a significant social and economic toll (Gore et al., 2011). Although these disorders emerge at developmentally distinct time points from early childhood in the case of autism (ASD) to young adulthood for schizophrenia (SZ), their genetic and environmental risk factors are surprisingly similar (Marin, 2016). Many of these risk factors affect the maternal-fetal environment and the most compelling of these insults is maternal infection. Indeed, epidemiological evidence indicates that maternal infection predisposes the developing fetus to a range of neurodevelopmental and psychiatric disorders (Brown and Patterson, 2011; Estes and McAllister, 2015; Knuesel et al., 2014; Patterson, 2002). Because numerous bacterial (Clarke et al., 2009; Sorensen et al., 2009), viral (Brown et al., 2004; Brown et al., 2001; Buka et al., 2008; Buka et al., 2001), and parasitic infections (Brown et al., 2005; Mortensen et al., 2007) are linked to increased risk of psychiatric illness, it appears to be the shared immune pathways activated in the mother in response to pathogen detection that confer risk to the fetus.

Rodent models of the environmental risk factor of infection during pregnancy (maternal immune activation or MIA) recapitulate many of the most replicated neuroanatomical, neurochemical, and functional features of ASD and SZ (Estes and McAllister, 2016). These features include reduced cortical thickness and increased ventricular volume (Li et al., 2009a; Piontkewitz et al., 2009), aberrations in Purkinje cells (Aavani et al., 2015; Shi et al., 2009), alterations in serotoninergic (Bitanihirwe et al., 2010; Reisinger et al., 2016; Winter et al., 2008) and dopaminergic signaling (Meyer et al., 2008; Ozawa et al., 2006; Vuillermot et al., 2010), deficits in the function of parvalbumin (PV) cells (Meyer et al., 2008; Shin Yim et al., 2017), and delay in the excitatory-to-inhibitory switch of GABA (Corradini et al., 2018) with the consequence of heightened excitatory neurotransmission and reduced long-range connectivity. These neurobiological alterations are accompanied by abnormal behaviors that resemble some of the core features of ASD and SZ such as abnormal communication and social interaction, repetitive behaviors, decreased sensorimotor gating, increased anxiety, deficits in working memory and cognitive flexibility and increased sensitivity to amphetamines (Estes and McAllister, 2016; Malkova et al., 2012; Meyer, 2014; Smith et al., 2007). While MIA likely acts as a primer for a range of neurodevelopmental disorders and is neither necessary nor sufficient at the population level to cause neuropsychiatric disorders (Estes and McAllister, 2016), the contribution of maternal infection to SZ alone may be as high as 30% of the population attributable risk (Brown and Patterson, 2011). Thus, disease mechanisms and biomarkers identified using MIA animal models, especially those focused on alterations in immune signaling in the brain, may have relevance for a substantial portion of affected individuals.

Changes in levels of maternal serum and fetal brain cytokines play a causal role in the MIA mouse model (Estes and McAllister, 2016). Maternal serum and placental interleukin-6 (IL-6) and interleukin-17a (IL-17a) increase exponentially during the acute phase of infection and are necessary and sufficient to cause ASD and SZ-relevant behaviors and neuropathologies in MIA offspring (Choi et al., 2016; Hsiao and Patterson, 2011; Smith et al., 2007). Alterations in fetal brain cytokine expression also occur within hours of infection, specifically upregulating canonical immune pathways (Arrode-Bruses and Bruses, 2012; Choi et al., 2016; Garbett et al., 2012; Meyer et al., 2006a), including increases in several pro-inflammatory cytokines (Fatemi et al., 2002; Meyer et al., 2008; Meyer et al., 2006b). Increases in maternal IL-17a cause elevations in fetal neuronal IL-17 receptor expression (Choi et al., 2016) and an increase in excitatory versus inhibitory connections in brain regions underlying repetitive and social behaviors (Shin Yim et al., 2017). Blocking the inflammatory cascade through anti-IL-6 or anti-IL-17 treatment in the maternal or fetal compartment prevents most MIA-induced behaviors and neuropathologies (Choi et al., 2016; Smith et al., 2007).

While the initial changes in cytokine signaling mediating the effects of MIA on fetal brain development are becoming more clear, it is unknown how these acute fetal cytokine signatures relate to changes in cytokine signaling in the brains of postnatal offspring that may correlate with the onset of clinically diagnosed behaviors that emerge in early life for ASD and young adulthood for SZ. Importantly, post-mortem studies of individuals with ASD and SZ show increases in some of the small number of mostly pro-inflammatory cytokines measured (Trepanier et al., 2016; Vargas et al., 2005). Using Luminex to measure protein levels of 23 cytokines from three brain regions and serum at 5 postnatal ages in the MIA mouse model, we similarly found that MIA leads to lifelong alterations in brain and serum cytokines in offspring (Garay et al., 2013). A wide range of cytokines were altered by MIA and each cytokine was altered in distinct age- and region-specific patterns that included decreases as well as increases, compared to controls. In frontal cortex (FC), cytokines were elevated at birth (postnatal day [P0]), decreased during the period of synapse formation and plasticity (P7-P30), followed by elevations again in early adulthood (P60). At birth, IL-1β, IL-10, IL-12 (p70), and GM-CSF were statistically significantly elevated while IL-1α, IL-4, and IL-6 trended toward higher levels. At P7, IL-2, IL-4, IL-5, IL-10, and IL-12 (p40) were decreased, while G-CSF was elevated. At P14 and P30, many cytokines were significantly decreased (P14: IL-1α, IL-1β, IL-2, IL-5, IL-9, IL-10, IL-13, Eotaxin, GM-CSF, IFNγ, and MCP-1; at P30: IL-1β, IL-3, IL-5, IL-6, IL-10, IL-12 (p40), IL-12 (p70), G-CSF, GM-CSF, MCP-1, and MIP-1β). Finally, in the adult, IL-1α, IL-6, IL-9, and IL-10 were elevated in the FC of MIA offspring compared to controls (Garay et al., 2013). Thus, MIA clearly leads to long-lasting changes in brain cytokine levels in offspring throughout postnatal development in a U-shaped age and region-dependent manner.

Since cytokines have known roles in the development, maturation, and plasticity of cortical connections (Deverman and Patterson, 2009), these observed decreases in cytokine levels during the period of synaptogenesis, dendritic spine formation, and activity-dependent plasticity may lead to chronic changes in cortical connectivity and altered behaviors in offspring. However, cytokines signal through specific cytokine receptors to alter cellular processes and it is levels of those receptors that are equally critical for signaling. Yet, it remains unknown whether MIA alters the expression of cytokine receptors in the brain of offspring throughout postnatal development. Here, we examined the expression levels of 21 cytokine receptors in the FC of MIA and control offspring using qPCR at the same five ages as our previous study (P0, P7, P14, P30 and P60) (Garay et al., 2013). We find that cytokine receptor expression is chronically altered in an age-specific manner, similar to the effects of MIA on cytokine levels in general over postnatal development. However, in contrast to the U-shaped change in cytokine protein levels over postnatal development (Garay et al., 2013), MIA-induced changes in cytokine receptor expression are more dynamic, exhibiting an oscillating pattern throughout postnatal development to adulthood. In FC of MIA offspring, the expression of a wide range of cytokine receptors is altered, with levels decreased at birth, dramatically increased at P7, decreased at P14, and increased to a lesser extent at P30 and P60. These alterations do not segregate easily into pro- or anti-inflammatory categories. Instead, they suggest chronic alterations in cytokine-mediated developmental processes. In early developmental ages, expression levels in cytokine receptors tend to change following MIA among individuals in a more tightly coordinated fashion than at later ages. Thus, in early development if MIA offspring differ in the expression of one cytokine receptor they are likely to differ in the expression of many. This pattern may reflect MIA-induced cytokine network feedforward and feedback regulation that is more pronounced in early development. Together with the U-shaped changes in cytokine protein levels during postnatal development (Garay et al., 2013), changes in cytokine receptor levels suggest an overall increase in cytokine signaling at P7, a dramatic decrease at P14 during periods of synapse formation, and an increase during periods of plasticity (P30) and early adulthood (P60).

## METHODS

### Animal Care and Use

All studies were conducted with approved protocols from the University of California, Davis Animal Care and Use Committee, in compliance with NIH guidelines for the care and use of experimental animals. For the MIA experiments C57BL/6 mice were bred in-house after purchase from Charles River. Mice were weaned at 21 days and housed with same-sex littermates.

### Maternal immune activation and tissue isolation

Pregnant C57BL/6 mice at gestational day (GD) 12.5 were injected intraperitoneally with fresh poly(I:C) dsRNA (Sigma Aldrich, St. Louis, MO, cat# P9582) at 20 mg/kg, or vehicle control (sterile .9% saline). Animals were weighed to confirm sickness-associated weight loss. Frontal cortex was dissected from 10 MIA and 10 saline male offspring at each age: P0, P7, P14, P30 and P60 and placed immediately in RNA*later* (Qiagen). Offspring were from at least 4 different litters for each age.

### RNA isolation and quantitative RT-PCR

RNA was isolated from mouse samples using RNeasy Mini Kit (Qiagen) and cDNA made using RT2 First Strand Kit (SA Biosciences). Custom 96-well RT2 Profiler PCR Arrays with 23 candidate genes and 2 controls were purchased from SA Biosciences. All samples were run in duplicate with a control sample on every plate. Samples were run on a Bio-Rad iCycler and analyzed using RT2 Profiler PCR Array Data Analysis version 3.5 (SA Biosciences). qPCR data were analyzed using the ddCt method with the housekeeping genes *heat shock protein 90* (*Hsp90*) and *glyceraldehyde 3-phosphate dehydrogenase phosphoglycerate* (*Gapdh*) as normalizers (Livak and Schmittgen, 2001).

### Statistical analysis

Data was subjected to two levels of significance: fold change and Wilcoxon-Mann-Whitney test with a Dunn-Šidàk correction for multiple comparisons. Candidate genes were chosen that showed greater than 2-fold change in comparison to controls and p ≤ 0.05. The fold changes were calculated based on age and sex-matched control offspring (i.e., non-MIA male-only offspring from animals injected with saline).

We subjected the log2 fold change expression results to principal components analysis (PCA). PCA is a dimension reduction technique which ordinates observations along uncorrelated, synthetically derived axes that maximize the variance each axis explains (Johnson and Wichern, 2007). The eigenvalues of the principal components (PCs) quantify the proportion of total variance each PC captures. We performed PCA in order to evaluate (a) whether individuals within an age group tend to cluster into a small number of gene expression subgroups, similar to the pattern observed for MIA outcomes, and (b) whether variation in gene expression is similarly concentrated for the different age groups. For each age group, we compared PC plots of the first two PCs to behavioral data to see if clustering patterns were similar. We also compared PC-to-PC eigenvalue decline among the different age samples to evaluate whether gene expression was similarly concentrated in a few major axes of variation, regardless of postnatal age. PCA was conducted with R statistical software (R Core Team, 2017) using base package functions.

## RESULTS

### MIA alters cytokine receptor profiles in FC in an age-specific manner

We first sought to characterize the expression of immune receptors using qPCR to complement our previous study examining the developmental profile of cytokine expression in mouse MIA offspring. A panel of 21 immune receptors (Table 1) was selected based upon the cytokines altered in our previous study (Garay et al., 2013) and those altered in a subset of individuals with ASD and SZ (Trepanier et al., 2016; Vargas et al., 2005). Receptors are grouped into functionally related subsets in Figure 1. Receptors for the classic pro-inflammatory cytokines responsible for early responses include *Il1r1* (*il-1 receptor type 1*), *Il6ra* (*il-6 receptor alpha*), *and Tnfr1* (*tumor necrosis factor receptor 1*). These receptors are grouped with the IL-1 family receptor *Il1rapl1* (*interleukin receptor accessory protein like-1*), *Ifngr1* (*interferon gamma receptor type 1*), *Ifitm3* (*interferon-induced transmembrane protein 3*), *and P2rx7* (*p2x purinoceptor 7)*, which can also mediate inflammatory innate responses. Additionally, the classic anti-inflammatory cytokine IL-10 keeps early inflammation in check and signals through *Il10ra* (*il-10 receptor alpha*). The next grouping of interleukin receptors mediate T-cell responses and include *Gmcsfr (granulocyte macrophage colony-stimulating factor receptor)*, which is functionally related to *Il3ra* (*interleukin 3 receptor alpha*) and *Il5ra* (*interleukin 5 receptor, alpha*). The next grouping contains chemokine receptors that attract immune cells to sites of inflammation; and, finally, three receptors involved in microglial function and the complement cascade. Five ages—P0, P7, P14, P30 and P60—were studied in the mouse model to parallel our previous study. These ages represent critical time points in brain development: namely, P0 is the period before most synapses have formed, P7 is the start of synaptogenesis, P14 is the peak of synaptogenesis, P30 is the peak of activity-dependent plasticity and P60 is early adulthood.

Principal component loadings for P1 and PC2.

Principal component loadings for P1 and PC2
| Genes | P0 |  | P7 |  | P14 |  | P30 |  | P60 |  |
| --- | --- | --- | --- | --- | --- | --- | --- | --- | --- | --- |
|  | PC1 | PC2 | PC1 | PC2 | PC1 | PC2 | PC1 | PC2 | PC1 | PC2 |
| Il2ra | 0.095086 | -0.36738 | NA | NA | 0.12285 | 0.01745 | -0.25348 | -0.3498 | -0.3037 | -0.2122 |
| Il9r | 0.204195 | -0.42048 | 0.51827 | -0.56683 | NA | NA | NA | NA | -0.53 | 0.51226 |
| Il5ra | NA | NA | 0.15757 | 0.02502 | 0.12172 | 0.10613 | 0.247933 | -0.2112 | 0.03793 | -0.1456 |
| Ccr2 | 0.151629 | -0.36598 | 0.32722 | -0.22639 | 0.2164 | 0.1218 | 0.546092 | 0.27705 | -0.2483 | 0.4046 |
| Il1rapl1 | -0.14918 | 0.0049 | -0.08814 | -0.09301 | 0.14341 | -0.16085 | 0.394392 | 0.03942 | -0.1717 | -0.0828 |
| Il3ra | -0.01805 | -0.29791 | 0.24413 | -0.3406 | 0.12377 | -0.06541 | 0.038966 | -0.0266 | -0.3621 | 0.10217 |
| Il17ra | -0.06397 | 0.24043 | -0.09341 | -0.13914 | 0.15198 | 0.45186 | 0.073906 | -0.1017 | 0.10928 | -0.1469 |
| Ccr5 | -0.09063 | -0.0573 | -0.14484 | 0.10708 | 0.34178 | 0.11041 | 0.061966 | 0.04347 | -0.1926 | -0.3428 |
| Ifitm3 | -0.0809 | -0.37159 | -0.15195 | -0.13475 | 0.25277 | 0.17027 | -0.01161 | 0.19489 | -0.1318 | -0.2606 |
| Cx3cr1 | -0.03402 | -0.25734 | -0.19728 | -0.27987 | 0.34026 | -0.16842 | 0.089925 | -0.3387 | -0.2379 | -0.1701 |
| Il6ra | -0.03169 | -0.03195 | -0.13648 | -0.11934 | -0.0132 | 0.63664 | 0.133747 | -0.4388 | -0.116 | 0.06978 |
| P2rx7 | -0.34846 | -0.03268 | -0.27628 | -0.13608 | 0.24162 | -0.04258 | 0.058191 | 0.09268 | 0.10821 | 0.02245 |
| Csf2ra | -0.25318 | 0.10705 | -0.35875 | -0.38809 | 0.24884 | -0.24601 | 0.380948 | 0.15245 | -0.1761 | 0.00732 |
| Ccr3 | -0.28157 | -0.28752 | -0.07221 | -0.05935 | 0.35935 | 0.20887 | 0.206005 | -0.1588 | -0.1644 | -0.1603 |
| Itgam | -0.21141 | -0.07646 | -0.08574 | -0.09138 | 0.28126 | -0.04162 | 0.194988 | -0.416 | -0.1337 | -0.1722 |
| Il10ra | -0.20997 | -0.07389 | -0.29515 | -0.11187 | 0.25128 | -0.1927 | 0.223199 | -0.2807 | -0.2506 | -0.3112 |
| Il1r1 | -0.2132 | -0.05442 | -0.0797 | -0.19003 | 0.2468 | 0.05726 | 0.278309 | -0.1042 | -0.0593 | -0.1517 |
| Cxcr2 | -0.22882 | 0.11919 | NA | NA | 0.02005 | 0.14893 | NA | NA | NA | NA |
| Tnfrsf1a | -0.2676 | 0.14332 | -0.20971 | -0.30769 | 0.18688 | -0.03765 | 0.129336 | 0.18537 | -0.2294 | -0.1043 |
| Cd55 | -0.55911 | -0.20145 | -0.10127 | -0.0405 | -0.004 | 0.25605 | 0.082291 | 0.0783 | -0.0181 | -0.1047 |
| Ifngr1 | -0.21108 | 0.08426 | -0.2338 | -0.18618 | 0.28219 | -0.16462 | -0.03261 | -0.1851 | -0.2447 | -0.2278 |

**Fig. 1.**
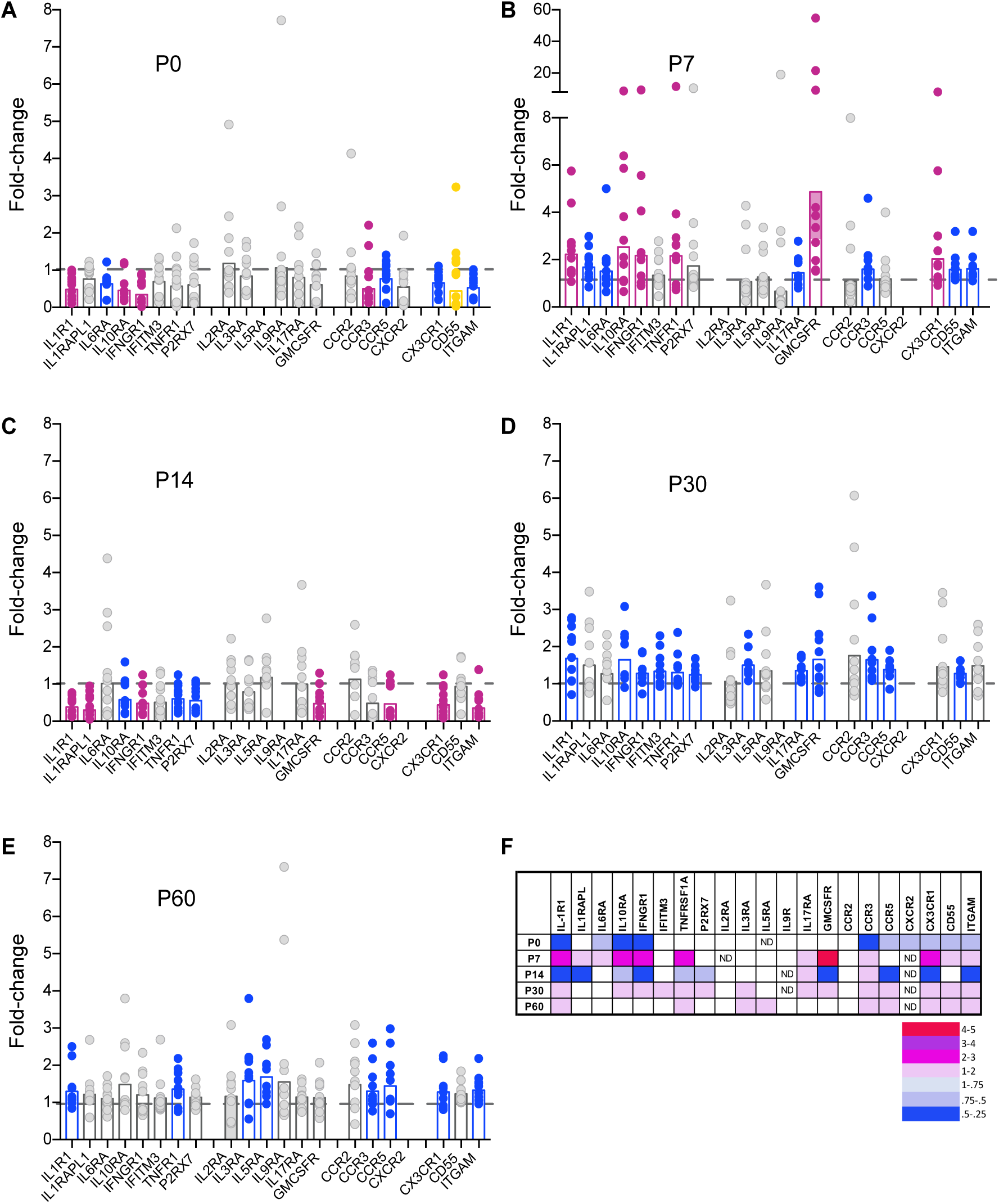
MIA induces long-lasting changes in the expression of cytokine receptors in frontal cortex throughout development. The fold-change in cytokine receptor expression in the FC of MIA offspring is plotted in comparison to saline offspring. Increases are shown above 1 and decreases, below. Statistically significant changes (p < 0.05) greater than 2-fold are represented in magenta, statistically significant changes less than 2-fold are represented in blue, and non-significant changes greater than 2-fold are represented in yellow. (A) At P0, 4 receptors were significantly decreased by at least 2-fold compared to controls and 4 more were significantly decreased less than 2-fold. CD55 was decreased more than 2-fold but was not statistically different from control values. (B) In contrast to the decreased levels at birth, 6 receptors were significantly <u>increased</u> at P7 by at least 2-fold and 6 more were significantly elevated at levels less than 2-fold. IL2RA and CXCR2 were below the level of detection. Please note the difference in scale of the graph in this panel. (C) At P14, 7 receptors were more than 2-fold decreased compared to controls. 3 additional receptors were significantly decreased compared to controls at levels less than 2-fold. IL9R and CXCR2 were below the level of detection. (D) At P30, 12 receptors were significantly increased compared to controls although none were increased above 2-fold. IL9RA and CXCR2 were below the level of detection. (E) At P60, 8 receptors were significantly elevated but less than 2-fold compared to controls. CXCR2 was below the level of detection. *n* = 10 brains from male offspring per treatment group from at least 4 independent litters. (F) The heatmap illustrates the changes in cytokine receptors over the 5 ages examined. Reds represent fold-increases and blues represent fold-decreases as indicated in the legend.

MIA causes long-lasting changes in expression of cytokine receptors from birth to early adulthood. Similar to MIA-induced changes in cytokines, the expression of cytokine receptors in MIA offspring was distinct at each age and altered in specific directions such that significantly altered cytokine receptors were either all increased or decreased at a given age when compared to controls (Figure 1). At birth and at P14, cytokine receptor levels were reduced relative to controls whereas receptor levels were elevated at P7, P30, and P60, showing an oscillating pattern of changes during postnatal development. At birth (Fig. 1A), 20 receptors were detectable and of these, 4 receptors were significantly decreased by at least 2-fold in the FC of MIA offspring compared to controls: *Ccr3* (*c-c motif chemokine receptor type 3*, 0.50-fold), *Il1r1* (0.48-fold), *Il10ra* (0.46-fold), and *Ifngr1* (0.34-fold). An additional 4 receptors were statistically significantly decreased compared to controls, although at levels less than 2-fold: *Il6ra* (0.64-fold), *Ccr5* (*c-c motif chemokine receptor type 5*, 0.77-fold), *Cx3cr1* (*cx3c chemokine receptor 1*, 0.69-fold), and *Itgam* (*integrin alpha m*, 0.53-fold). Finally, one receptor—*Cd55* (*complement decay-accelerating factor*, 0.44-fold)—was decreased more than 2-fold but was not statistically different from control values.

In contrast to the decreased levels at birth, six cytokine receptors were significantly <u>increased</u> at P7 by at least 2-fold in the FC of MIA offspring compared to controls (Fig. 1B): *Il1r1* (2.23-fold), *Il10ra* (2.5-fold), *Ifngr1* (2.18-fold), *Cx3cr1* (2.03-fold), *Tnfr1* (2.16-fold) and *Gmcsfr* (4.87-fold). All of these receptors are significantly decreased either at birth or P14 with the exception of *Tnfr1*. An additional 6 receptors were significantly increased compared to controls at levels less than 2-fold: *Il1rapl1* (1.69-fold), *Il6ra* (1.52-fold), *Il17ra* (*interleukin receptor 17, alpha*, 1.44-fold), *Ccr3* (1.60-fold), *Cd55* (1.59-fold), and *Itgam* (1.61-fold).

Cytokine receptor expression changed direction again at P14 with 7 out of the 19 detectable receptors significantly <u>decreased</u> more than 2-fold in the FC of MIA offspring compared to control (Fig. 1C). Two of the cytokine receptors decreased at P0 were also significantly decreased at P14: *Il1r1* (0.39-fold), and *Ifngr1* (0.48-fold) along with *Il1rapl1* (0.30-fold), *Ccr5* (0.47-fold), *Cx3cr1* (0.44-fold), *Itgam* (0.36-fold), and *Gmcsfr* (0.48-fold). An additional 3 receptors were significantly decreased compared to controls, although at levels less than 2-fold: *Il10ra* (0.58-fold), *Tnfr1* (0.60-fold), and *P2rx7* (0.56-fold).

At older ages, the direction of change in cytokine receptors reversed again. While no receptors were more than 2-fold changed in the FC of MIA offspring at the older ages of P30 and P60, many cytokine receptors were significantly <u>increased</u> at lower levels compared to controls at these ages. At P30, 12 out of 19 receptors were significantly elevated over controls: *Il1r1* (1.68-fold), *Il3ra* (1.50-fold), *Ccr3* (1.65-fold), *Il10ra* (1.65-fold) *Ifngr1* (1.28-fold), *Ifitm3* (1.33-fold), *Tnfr1* (1.31-fold), *P2rx7* (1.24-fold), *Il17ra* (1.36-fold), *Gmcsfr* (1.66-fold), *Ccr5* (1.39-fold), and *Cd55* (1.28-fold) (Fig. 1D). In adults (Fig. 1E), 8 of the 20 detectable cytokine receptors were elevated in the FC of MIA offspring compared to controls. *Il3ra* (1.59-fold) and *Il5ra* (1.69-fold) were significantly increased by at least 1.5-fold and several additional receptors were elevated to a lesser extent: *Il1r1* (1.30-fold), *Tnfr1* (1.36-fold), *Ccr3* (1.31-fold), *Ccr5* (1.44-fold), *Cx3cr1* (1.28-fold), and *Itgam* (1.33-fold).

Of the many cytokine receptors altered in the FC of MIA offspring, five were significantly changed at multiple ages at levels over 2-fold relative to controls—*Il1r1, Ifngr1, Il10ra, Cx3cr1 and Gmcsfr*—suggesting persistent dysfunction within those pathways (Fig. 1). At birth, three receptors met the criteria and were significantly lower than controls: *Il1r1* (0.48-fold), *Il10ra* (0.46-fold), and *Ifngr1* (0.34-fold). All three of these receptors were significantly increased by P7 (*Il1r1* (2.23-fold), *Il10ra* (2.53-fold) and *Ifngr1* (2.18-fold)) as well as two other receptors: *Cx3cr1* (2.03-fold) and *Gmcsfr* (4.87-fold). At P14, all but one of these receptors was significantly decreased (*Cx3cr1* (0.44-fold), *Gmcsfr* (0.48-fold), *Il1r1* (0.39-fold), and *Ifngr1* (0.48-fold)). Thus, of the five targets, three receptors (*Il10ra, Cx3cr1*, and *Gmcsfr*) were significantly altered at two ages while two receptors (*Il1r1* and *Ifngr1*) were significantly altered at multiple ages in an oscillating pattern.

The greatest magnitude of change in cytokine receptor expression occurred at P7 when the statistically significant fold changes ranged from 1.4 to 4.8 compared to ranges of 0.34-1.68 at the other ages. The earliest 3 ages were also the times when the largest number of different cytokines were altered above a 2-fold threshold. For cytokine receptors that were statistically different from controls, the following percentages of total detectable receptors were altered: P0: 45%, P7: 63%, P14: 53%, P30: 63%, and P60: 40%. For cytokine receptors that were both significantly different at least 2-fold, the changes were: P0: 20%, P7: 31%, P14: 37%, P30: 0%, P60: 0%. Not all receptors were detected with these primers at every age. Four receptors were below the level of detection of this assay in an age-specific manner: specifically, *Il2ra, Il5ra, Il9ra*, and *Cxcr2*. *Il2ra* and *Il5ra* were undetectable at P0 and P7; *Il9ra* at P14 and P30; and *Cxcr2* at every age except for P0. Together, these results indicate that, like cytokines (Garay et al., 2013), cytokine receptors are altered in the FC of MIA offspring in an age-specific, cyclical developmental pattern. Additionally, the subset of cytokine receptors altered and the direction of change are age-dependent, with the largest changes occurring in early development at P7 and P14, the period of peak synaptogenesis and spine formation.

### MIA offspring show different development patterns of variance in gene expression

Behavioral and neuroanatomical outcomes in MIA offspring can be highly variable within and between litters as well as between studies (Estes and McAllister, 2016). This heterogeneity is also reflected in cytokine receptor expression, with some individuals showing modest differences from controls and others displaying large changes across numerous genes. We therefore tested: (1) whether cytokine receptor expression patterns segregated MIA offspring into two populations (similar to what is observed in studies of MIA outcomes where some offspring are not affected), and (2) whether expression in a small subset of cytokine receptors accounted for most variance between individuals.

To evaluate age-group variability and clustering, we performed PCA for the multivariate gene expression panel. Within an age group, there were occasional outlier individuals, but no apparent tendency for MIA offspring to cluster into gene expression subgroups on the two largest PCs (Supplemental Figure 1). Thus, while some MIA offspring can appear grossly normal in behavioral assays, this subset is not clearly identifiable through examination of cytokine receptor expression changes, which appear more continuous.

To evaluate whether expression in a subset of cytokine receptors tends to define the major axes of variation, or if instead variation on these axes is distributed more evenly over the gene assay, the minimum gene subset needed to account for 80% of variance explained by the first two PCs was determined. In every age group, the 80% threshold required at least 10 genes (Supplemental Table 1), indicating a fairly even distribution. To gain greater clarity on whether variation in gene expression tends to be concentrated in just a few PCs, we plotted eigenvalue decay for the age class PCAs. While PCs 1 and 2 account for the majority of gene expression variance in all age classes, the cumulative proportion is substantially higher in P0 and P7 (83% of variance) than in P14, P30 or P60 (60-68%) (Fig. 2; Supplemental Figure 1). This indicates relatively more correlated gene expression at early postnatal ages, which becomes more heterogeneous over post-natal development. These results are also consistent with other gene expression studies which find only modest MIA-induced changes in adult cortex despite the presence of disease-relevant neuropathologies and behaviors (Connor et al., 2012; Smith et al., 2007). The shared genes driving variance at birth and at P7 may be associated with susceptibility and/or resilience to MIA offspring outcomes. Taken together, these data suggest that MIA has the greatest impact on cytokine receptor expression early in development.

**Fig. 2.**
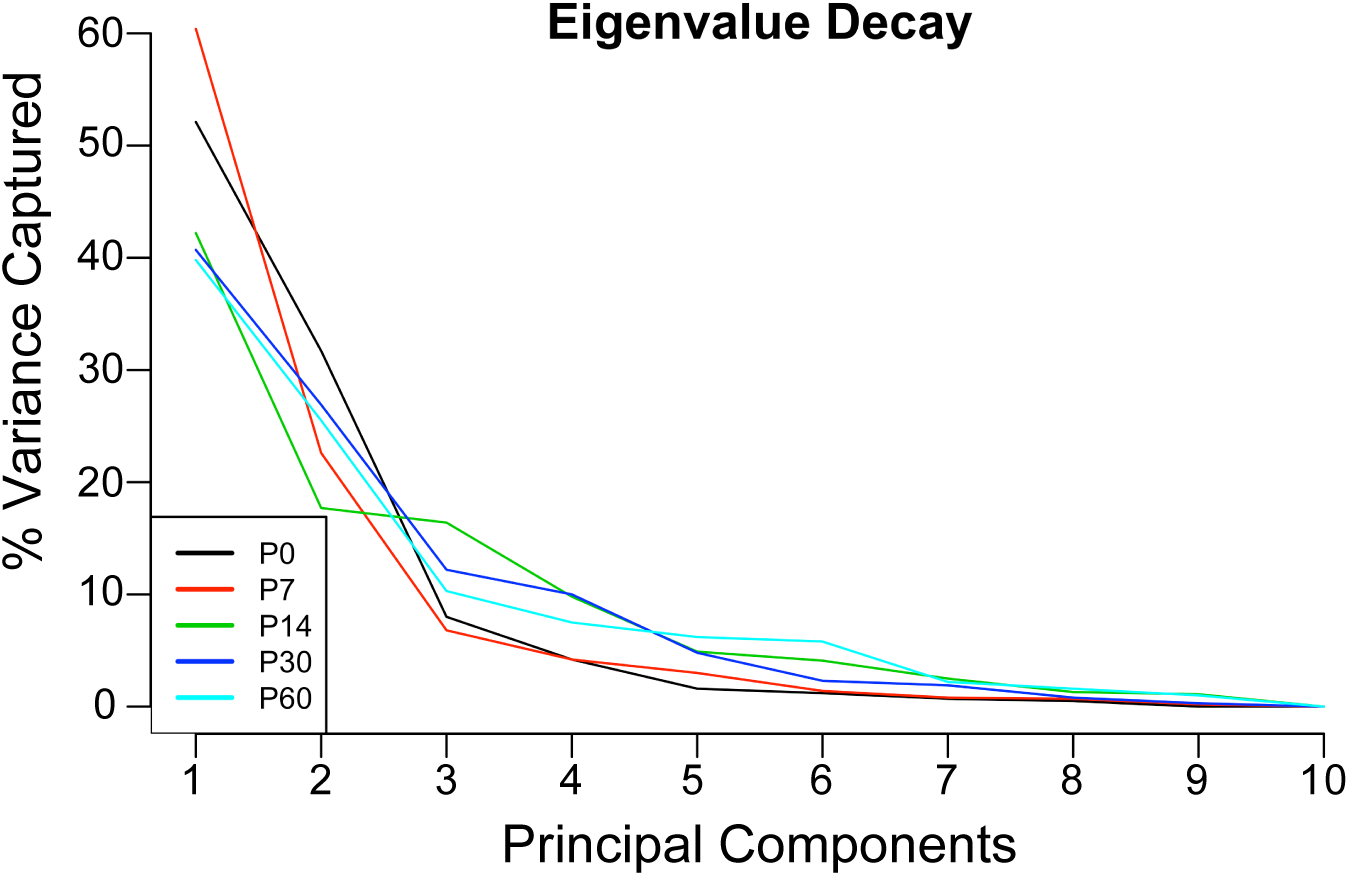
MIA offspring show a shift in cytokine receptor expression patterns during synaptogenesis. Scree plot representing the percent of variance accounted for by the principal components (PCs) at each age. The concentration of cytokine receptor expression variation clusters into two groups: P0-P7 and P14-P60. At P0 and P7, variation is concentrated in the first two PCs. At later ages, the variance is more evenly dispersed across the PCs.

## Discussion

In this study, the effect of poly(I:C)-induced MIA on cytokine receptor expression in the FC of offspring from birth to adulthood was examined. In concordance with our previous study on cytokine protein levels (Garay et al., 2013), cytokine receptor expression is significantly altered in an age-specific manner in offspring, with all significant changes following the same direction at a particular age. Across ages the direction of change of cytokine receptors in MIA offspring shows an oscillating pattern—down at birth, up at P7, down at P14, and up at later ages—as compared to control offspring. These changes do not segregate easily into pro- or anti-inflammatory categories and instead indicate chronic alterations in cytokine-mediated developmental processes, including aberrant microglial function (though not inflammatory), neuronal differentiation, synapse formation and elimination, and synaptic function, that could contribute to the ASD- and SZ-like neuropathology and behavioral alterations in MIA offspring.

Growing evidence implicates lifelong immune alterations in individuals with ASD and SZ. Changes in peripheral immune markers are associated with both disorders but study results have been mixed and peripheral immune markers have not shown the specificity, reliability, and positive predictive value to be of use in a clinical setting. Cytokines are also altered in postmortem brain tissue from ASD and SZ individuals (Estes and McAllister, 2015; Trepanier et al., 2016; Vargas et al., 2005), although these studies are limited by the typically small number of mostly pro-inflammatory cytokines measured. The increased expression of a few proinflammatory cytokines in brain tissue has led to the hypothesis that chronic neuroinflammation drives the clinical course of ASD and SZ. However, chronic neuroinflammation cannot be assessed from the mere presence or absence of cytokines without the parallel presence of other signs of neuroinflammation such as breaches in the blood-brain-barrier (BBB), gliosis, and peripheral immune cell infiltration (Estes and McAllister, 2014). Importantly, these definitive criteria for neuroinflammation do not appear to be present in humans or in MIA rodent models (Borta and Schwarting, 2005; Collste et al., 2017; Cotel et al., 2015)(Giovanoli et al., 2016; Giovanoli et al., 2015; Holmes et al., 2016). In particular, our previous discovery of the age-dependent changes in cytokines in the FC of MIA offspring throughout postnatal development does not support the neuroinflammation hypothesis. Further, we found no evidence of decreased BBB integrity, immune cell infiltration or gliosis in the brains of MIA offspring (Garay et al., 2013). Finally, the oscillating pattern of changes in cytokine receptor expression across postnatal development reported here also does not support a clear role for neuroinflammation in offspring caused by MIA. Together, these results from humans and animal models indicate that it is the dynamic dysregulation of immune molecule signaling in the brain, rather than neuroinflammation, that contributes to ASD and SZ-related neuropathology.

The oscillating changes in cytokine receptors in the FC of offspring throughout development likely arise from complex feedforward and feedback loops that cytokine networks exert on cytokine and receptor expression as a means of maintaining homeostasis. For example, cytokines often regulate the expression of their receptors and others. Thus, an increase in the production of a cytokine may be met with a parallel decrease in cytokine receptor expression bringing the overall level of signaling back to physiological levels. In other contexts, increases or decreases in cytokine signaling could result in a similar directional change in receptor expression leading to an amplification of the effect. When specific antibodies to cytokine receptors are developed in the future, assessing cytokine receptor protein levels will provide important insight into the overall change in cytokine signaling in the brains of offspring following MIA that is predicted based on mRNA expression levels. Moreover, understanding the molecular mechanisms that mediate the effects of MIA in causing this oscillating pattern of changes in cytokine receptors is a major goal for future studies.

While changes in prenatal cytokines have been shown to be causal for some MIA-induced behaviors (Choi et al., 2016; Shin Yim et al., 2017; Smith et al., 2007), the presence and effects of long-lasting changes in cytokine signaling throughout postnatal development are largely unknown. In the context of infection, cytokines are classified as having mostly pro or anti-inflammatory activity, but under physiological conditions they play diverse roles in brain development and maintenance (Deverman and Patterson, 2009). In early development, many cytokines act as patterning and neurotrophic factors and in postnatal development, cytokines promote synapse formation and elimination, modulate synaptic strength, and alter the release of numerous neurotransmitters and hormones from neurons and glia (Garay and McAllister, 2010)(Deverman and Patterson, 2009). The MIA-induced chronic alterations in cytokines and their receptors could represent homeostatic, adaptive or pathological processes with or without an immune etiology in an age-dependent manner. These physiological cytokine-mediated developmental and homeostatic processes must be altered by MIA-induced changes in cytokine signaling in the brains of offspring.

A surprisingly large number of cytokine receptors were altered at each age postnatally in the FC of offspring following MIA. Because so many cytokine receptors were altered in distinct patterns at each age, future work is essential for determining which, if any, of the altered cytokine receptors could be driving changes in the other receptors and which are critical for changes in cortical connectivity and/or function in offspring at each developmental age. PCA analysis revealed a developmental shift in relatively more correlated cytokine receptor expression at early postnatal ages (P0-P7), which becomes more heterogeneous over post-natal development. Future work is needed to determine if this represents a physiological property of cytokine networks or an MIA-induced shift in the coordinated activity of cytokine receptor expression. It will also be important to identify shared downstream signaling hubs, like JAK/STAT and major histocompatibility complex I (MHCI) molecules, that may translate changes in the many cytokine receptors on each cell into overall changes in neuronal or glial function.

One approach to identifying the most important cytokine receptors altered by MIA is to focus on those that are significantly changed at multiple ages at levels over 2-fold relative to controls, suggesting persistent dysfunction within those pathways. Five cytokine receptors met these criteria: *Il1r1, Ifngr1, Il10ra, Cx3cr1* and *Gmcsfr*. Each of these receptors are discussed in depth below. These chronically altered cytokine receptors may provide the most insight into the causal immune signaling pathways underlying neuropathology and the emergence of disease-related behaviors. Importantly, they may also have the greatest potential as diagnostic biomarkers or as targets for development of new therapeutics.

Receptors for two cytokine families—IL1 and IFNγ—that are generally considered pro-inflammatory were altered throughout development in the FC of MIA offspring. Expression of *Il1r1* was significantly decreased at birth, increased at the start of synaptogenesis, decreased during the peak of synaptogenesis and spine formation, and increased below 2-fold during the peak of activity-dependent plasticity and early adulthood. IL-1R1 is the canonical IL-1 receptor, binds the pro-inflammatory cytokines IL-1α and β, and is expressed on both neurons and glia throughout the brain. Aside from its inflammatory properties, IL-1R1 signaling plays roles in neurogenesis, neuronal differentiation, plasticity, synapse formation, and neuroexcitotoxicity (Yirmiya and Goshen, 2011). Another member of the IL-1 family, *Il1rapl1* was significantly increased at P7 (less than 2-fold) during the start of synaptogenesis and decreased more than 2-fold at P14, during the peak of synaptogenesis and spine formation. *IL1RAPL1* was identified in a screen for X-linked intellectual disability and is similar to the IL-1 accessory proteins (Bhat et al., 2008). Intriguingly, IL1RAPL1 acts as a synaptic organizer and is a genetic risk factor for ASD and SZ (Bhat et al., 2008; Melhem et al., 2011; Valnegri et al., 2011). Mutations in *IL1RAPL1* associated with neurodevelopmental disorders lead to reduced synapse formation (Pavlowsky et al., 2010; Valnegri et al., 2011). An MIA-induced reduction in *Il1rapl1* expression during periods of synapse formation may replicate the effect of this genetic risk factor leading to altered connectivity.

The pro-inflammatory cytokine receptor, *Ifngr1*, exhibited similar changes in expression following MIA throughout cortical development as *Il1r1* although it was not altered by early adulthood. IFN-γR1 is the ligand-binding chain (alpha) of the heterodimeric interferon gamma receptor, which mediates the signaling of IFN-γ. IFN-γ signaling during brain development promotes neurite outgrowth and neuronal survival (Song et al., 2005) and may also impact synapse formation. IFN-γ induces surface major histocompatibility complex I (sMHCI) expression in neurons, which regulates synapse formation and function (Elmer et al., 2013; Glynn et al., 2011)(Neumann et al., 1997; Victorio et al., 2012). Indeed, we have previously shown that MIA causes a deficit in the ability of cortical neurons to form synapses that is dependent on a MHCI and myocyte enhancer factor-2 (MEF2) signaling pathway (Elmer et al., 2013). Alterations in IFN-γ signaling may contribute to this aberrant brain connectivity by regulating sMHCI. IFN-γ signaling has also been implicated in excitoneurotoxicity. Specifically, association between IFN-γR1 and the AMPAR subunit GluR1 causes IFN-γ-mediated phosphorylation of GluR1 and enhanced Ca2+ influx through calcium permeable AMPARs, which in turn leads to dendritic beading and cell death (Mizuno et al., 2008). Thus, changes in cortical connectivity and cell number may result from MIA-induced alterations in IFNGR1 expression.

A prototypical anti-inflammatory receptor—*Il10ra*—was also altered in the FC of offspring in a similar pattern as *Il1r1* and *Ifngr1*. *Il10ra* expression was decreased at birth and increased at the start of synaptogenesis over 2-fold, and also decreased during the peak of synaptogenesis and spine formation and increased during the peak of activity-dependent plasticity below 2-fold. As a prototypical anti-inflammatory cytokine in the periphery, IL-10 prevents excessive, unresolved inflammation and limits secondary tissue damage. Thus, it is unsurprising to see alterations in IL-10 signaling in parallel with the prototypical inflammatory cytokines IL-1β and IFN-γ. In the brain, IL-10 has been reported to regulate neurite outgrowth and synapse formation following oxygen-glucose deprivation-induced injury (Chen et al., 2016). IL-10 signaling through JAK/STAT upregulates Netrin-1, which is essential for axon regeneration in this model and suggests that IL-10-induced expression of Netrin-1 may occur under physiological conditions where Netrin-1 plays essential roles in neuronal development, and axonal growth and branching.

Several receptors implicated in microglial function and microglia-neuron communication were also altered more than 2-fold in the FC of MIA offspring at multiple ages examined. *Cx3cr1* and *Gmcsfr* levels were most strongly increased at the beginning of synaptogenesis (P7) and decreased during the peak of synaptogenesis and spine formation (P14). Along with two other receptors significantly decreased at P14 (*Itgam* and *Ccr5*), these receptors implicate microglial dysfunction in MIA-induced changes in cortical connectivity. Intriguingly, this age coincides with a critical developmental window when microglia sculpt forming circuits (Prinz and Priller, 2014).

CX3CR1, also known as the fractalkine receptor or G-protein coupled receptor 13 (GPR13), is required for microglia-mediated synaptic pruning in early development and spine formation and elimination in mature circuits (Paolicelli et al., 2011; Zhan et al., 2014). CX3CR1 binds the chemokine CX3CL1 (also called neurotactin or fractalkine), which is expressed in neurons and mediates the recruitment of CX3CR1-expressing microglia to neurons following injury (Paolicelli et al., 2011) and to synapses during activity-dependent synapse elimination (Schafer et al., 2013). Local neuron-microglia signaling through CX3CR1-CX3CL1 has been proposed to serve as an instructive signal for complement-mediated synaptic pruning (Schafer et al., 2013). Like CX3CR1, CR3, also known as macrophage-1 antigen or integrin αM (ITGAM), is also expressed on microglia. CR3 is a complement receptor consisting of CD11b (integrin αM) and CD18 (integrin β2). CR3 is found on several types of immune cells and binds to C3b and C4b, which aid in complement-mediated opsonization of bacteria and debris. Microglial-expressed CR3 in the brain recognizes tagged synapses and initiates synaptic pruning at a subset of synapses (Schafer et al., 2012). Taken together, alterations in CR3 and CX3CR1 expression seen in the FC of MIA offspring suggest age-specific changes in microglial migration, complement-mediated synaptic pruning, and adaptive plasticity.

Signaling through GMCSFR in the periphery promotes microglial proliferation (Lee et al., 1994) and is essential for maturation into antigen-presenting cells (Re et al., 2002). Although little is known about the role for GM-CSF in postnatal brain development, it has been reported that GMCSFR is expressed on neurons (Schabitz et al., 2008) and can stimulate proliferation and inhibit apoptosis of neural progenitor cells (Kim et al., 2004; Kim et al., 2009). GMCSF is also essential for spine formation in hippocampal neurons and for hippocampal-dependent spatial and fear memory in young adult mice (Krieger et al., 2012). Finally, CCR5 binds the chemokines RANTES, MIP, and CCL3L1 and its role in the brain is not fully known. In the periphery, CCR5 is expressed on microglia and alters microglial activation and function (Maung et al., 2014). Recently, like GMCSFR, CCR5 has also been shown to be expressed on astrocytes and neurons in the central nervous system, where CCR5 negatively regulates synaptic plasticity and hippocampal learning and memory (Zhou et al., 2016).

Although these changes in cytokine receptors suggest functional sequelae during FC development, it is important to note that changes in expression of each receptor are often distinct from changes in the protein levels of their corresponding cytokines, as detected in our previous study (Garay et al., 2013). Cytokine protein levels are generally elevated at birth, decreased from P7-P14-P30, and then elevated again in early adulthood (P60). Together with this U-shaped trajectory of cytokine protein levels during postnatal development (Garay et al., 2013), the oscillating changes in cytokine receptor levels reported here suggest dynamic changes in connectivity with age. At birth, cytokines are elevated while cytokine receptors are lower making it difficult to predict any effect on overall cytokine signaling. At P7, cytokine levels are lower, but their receptor expression is much higher, suggesting a compensatory effect and possibly an overall increase in cytokine signaling at P7. At P14, both cytokines and their receptors are decreased indicating that cytokine signaling is most likely dramatically decreased during this period of peak synaptogenesis and spine formation. At P30, changes in cytokines again oppose the direction of change in their receptors, making it difficult to infer what the overall change in cytokine signaling might be. However, at P60, both cytokines and their receptors are elevated indicating a clear increase in cytokine signaling.

Despite tremendous growth in our understanding of the genetic architecture and biological underpinnings of psychiatric disorders, there have been no new classes of drugs developed in over fifty years (Agid et al., 2007; Insel, 2012; Millan et al., 2016). Current pharmaceuticals developed for psychiatric disorders were reverse engineered from clinical observation of repurposed drugs with psychoactive properties; no drug has yet been created based upon the neurobiological substrates of these disorders (Agid et al., 2007). There are many reasons for our failure to identify new molecular targets. Most preclusive, diagnosis relies upon descriptive psychopathology, leading to patient cohorts representing a wide range of etiology, neuropathology, clinical course, and response to treatment (Marin, 2016; Millan et al., 2016). Drug discovery would benefit greatly from the stratification of patients—ideally, based upon a biological marker—into etiologically homogeneous subgroups. A key advance for the field would be identification of biomarkers that identify at-risk individuals prior to full clinical manifestation. Biomarkers that aid in early identification and intervention could be transformative for individuals predisposed to these disorders even in the absence of a complete mechanistic understanding or cure (Abi-Dargham and Horga, 2016). Most beneficial to this discovery are animal models in which neuroanatomical, functional, and behavioral abnormalities emerge at developmentally specific time points paralleling the alterations in developmental trajectories found in these disorders (Estes and McAllister, 2016). In these animal models, disease course and developmental trajectory can be interrogated, which may assist in the identification of therapeutic windows and biomarkers preceding full conversion from prodromal states.

Studies examining the progression of the disease process through changes in expression of immune molecules over time during brain development in disease models, such as the present study, are critical for determining the timing of the onset of pathology, the biomarkers that distinguish different phases of disease progression, and novel therapeutic targets specific for distinct stages of the disease process. This approach is clearly important for diseases initiated by maternal infection, such as ASD and SZ, since the changes in immune signaling in the brain are so dramatically different across postnatal development. For example, decreasing cytokine signaling through anti-inflammatory therapies could alleviate the effects of elevated cytokine signaling in early adulthood, but this approach would likely exacerbate pathology when cytokine signaling is decreased during earlier stages of synaptogenesis. Identifying novel ligands for molecular imaging studies that could signal the stage of the disease process (increased or decreased immune molecule signaling) in vivo, would increase the success of targeted treatments. Altered brain cytokines may meet these needs, but identification of brain biomarkers in vivo requires molecular imaging (Abi-Dargham and Horga, 2016). Most cytokines are soluble proteins with short half-lives, which makes them poor candidates for molecular imaging in vivo using positron emission tomography (PET). Cytokine receptors are more suitable as ligands for generating radiotracers. The receptors discovered in this study provide several novel targets for such biomarkers and imaging approaches. Unlike peripheral immune measures, levels of brain cytokines may be a direct readout of ongoing pathophysiology relevant to ASD and SZ diagnosis (Fillman et al., 2013; Li et al., 2009b; Trepanier et al., 2016; Vargas et al., 2005; Wei et al., 2011). Similarly, small molecules targeting these specific altered immune signaling pathways in the brain are potentially exciting new therapeutics for a range of neuro-immune based CNS disorders. Developing these biomarkers and therapeutics is critical to fill the current lack of any viable approach to image immune signaling in vivo, potentially transforming diagnostics and treatment of a wide range of neuropsychiatric disorders.

---

**Supplemental Figure 1.**
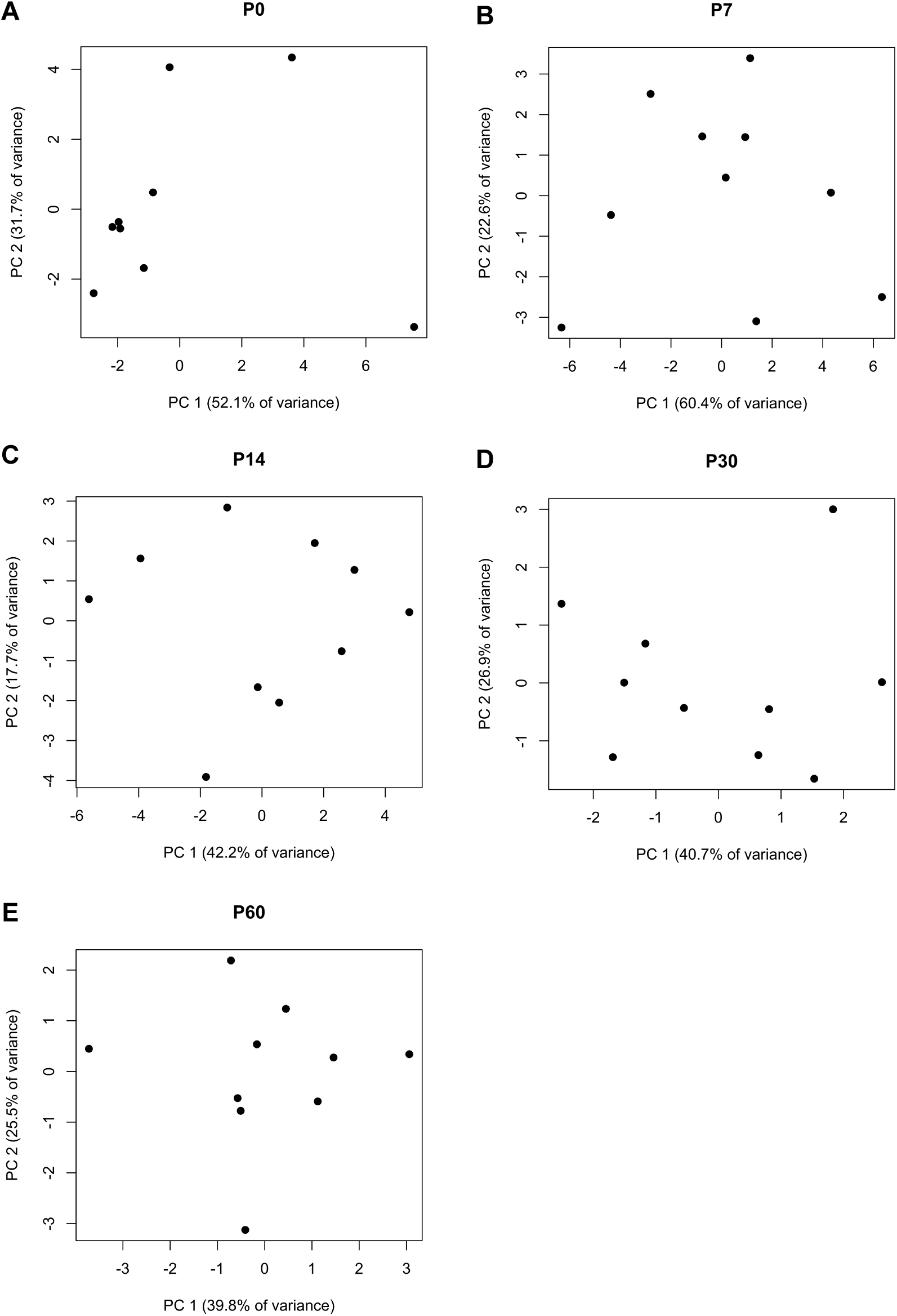
Scatterplot of specimen PC scores on the first and second PCs, obtained from PCA analyses of MIA offspring at each age. (A-E) The distribution of specimens reflects how animals differ from one another along the primary axes of gene expression variation. PCA analyses do not reveal a grouping structure within age samples.

## Acknowledgments

We wish to thank Dr. David C. Katz for consultation on statistics and editing the manuscript. This work was supported by a Stanley & Jacqueline Schilling Neuroscience Postdoctoral Research Fellowship (M.L.E.), Dennis Weatherstone Predoctoral Fellowships from Autism Speaks (#7825 M.L.E. and #6339 B.M.E.), the Letty and James Callinan and Cathy and Andrew Moley Fellowship from the ARCS Foundation (M.L.E.), a Dissertation Year Fellowship from the University of California Office of the President (M.L.E.), P50-MH106438-01 (C.S.C), and the University of California Davis Research Investments in Science and Engineering Program (A.K.M.).

## References

Aavani, T., Rana, S.A., Hawkes, R., Pittman, Q.J., 2015. Maternal immune activation produces cerebellar hyperplasia and alterations in motor and social behaviors in male and female mice. Cerebellum 14, 491–505.

Abi-Dargham, A., Horga, G., 2016. The search for imaging biomarkers in psychiatric disorders. Nat Med 22, 1248–1255.

Agid, Y., Buzsaki, G., Diamond, D.M., Frackowiak, R., Giedd, J., Girault, J.A., Grace, A., Lambert, J.J., Manji, H., Mayberg, H., Popoli, M., Prochiantz, A., Richter-Levin, G., Somogyi, P., Spedding, M., Svenningsson, P., Weinberger, D., 2007. How can drug discovery for psychiatric disorders be improved? Nat Rev Drug Discov 6, 189–201.

Arrode-Bruses, G., Bruses, J.L., 2012. Maternal immune activation by poly I:C induces expression of cytokines IL-1beta and IL-13, chemokine MCP-1 and colony stimulating factor VEGF in fetal mouse brain. Journal of neuroinflammation 9, 83.

Bhat, S.S., Ladd, S., Grass, F., Spence, J.E., Brasington, C.K., Simensen, R.J., Schwartz, C.E., Dupont, B.R., Stevenson, R.E., Srivastava, A.K., 2008. Disruption of the IL1RAPL1 gene associated with a pericentromeric inversion of the X chromosome in a patient with mental retardation and autism. Clin Genet 73, 94–96.

Bitanihirwe, B.K., Peleg-Raibstein, D., Mouttet, F., Feldon, J., Meyer, U., 2010. Late prenatal immune activation in mice leads to behavioral and neurochemical abnormalities relevant to the negative symptoms of schizophrenia. Neuropsychopharmacology 35, 2462–2478.

Borta, A., Schwarting, R.K., 2005. Post-trial treatment with the nicotinic agonist metanicotine: Differential effects in Wistar rats with high versus low rearing activity. Pharmacol Biochem Behav 80, 541–548.

Brown, A.S., Begg, M.D., Gravenstein, S., Schaefer, C.A., Wyatt, R.J., Bresnahan, M., Babulas, V.P., Susser, E.S., 2004. Serologic evidence of prenatal influenza in the etiology of schizophrenia. Arch Gen Psychiatry 61, 774–780.

Brown, A.S., Cohen, P., Harkavy-Friedman, J., Babulas, V., Malaspina, D., Gorman, J.M., Susser, E.S., 2001. A.E. Bennett Research Award. Prenatal rubella, premorbid abnormalities, and adult schizophrenia. Biological psychiatry 49, 473–486.

Brown, A.S., Patterson, P.H., 2011. Maternal infection and schizophrenia: implications for prevention. Schizophr Bull 37, 284–290.

Brown, A.S., Schaefer, C.A., Quesenberry, C.P., Jr., Liu, L., Babulas, V.P., Susser, E.S., 2005. Maternal exposure to toxoplasmosis and risk of schizophrenia in adult offspring. Am J Psychiatry 162, 767–773.

Buka, S.L., Cannon, T.D., Torrey, E.F., Yolken, R.H., Collaborative Study Group on the Perinatal Origins of Severe Psychiatric, D., 2008. Maternal exposure to herpes simplex virus and risk of psychosis among adult offspring. Biological psychiatry 63, 809–815.

Buka, S.L., Tsuang, M.T., Torrey, E.F., Klebanoff, M.A., Wagner, R.L., Yolken, R.H., 2001. Maternal cytokine levels during pregnancy and adult psychosis. Brain Behav Immun 15, 411–420.

Chen, H., Lin, W., Zhang, Y., Lin, L., Chen, J., Zeng, Y., Zheng, M., Zhuang, Z., Du, H., Chen, R., Liu, N., 2016. IL-10 Promotes Neurite Outgrowth and Synapse Formation in Cultured Cortical Neurons after the Oxygen-Glucose Deprivation via JAK1/STAT3 Pathway. Sci Rep 6, 30459.

Choi, G.B., Yim, Y.S., Wong, H., Kim, S., Kim, H., Kim, S.V., Hoeffer, C.A., Littman, D.R., Huh, J.R., 2016. The maternal interleukin-17a pathway in mice promotes autism-like phenotypes in offspring. Science 351, 933–939.

Clarke, M.C., Tanskanen, A., Huttunen, M., Whittaker, J.C., Cannon, M., 2009. Evidence for an interaction between familial liability and prenatal exposure to infection in the causation of schizophrenia. Am J Psychiatry 166, 1025–1030.

Collste, K., Plaven-Sigray, P., Fatouros-Bergman, H., Victorsson, P., Schain, M., Forsberg, A., Amini, N., Aeinehband, S., Karolinska Schizophrenia Project, c., Erhardt, S., Halldin, C., Flyckt, L., Farde, L., Cervenka, S., 2017. Lower levels of the glial cell marker TSPO in drug-naive first-episode psychosis patients as measured using PET and [(11)C]PBR28. Mol Psychiatry 22, 850–856.

Connor, C.M., Dincer, A., Straubhaar, J., Galler, J.R., Houston, I.B., Akbarian, S., 2012. Maternal immune activation alters behavior in adult offspring, with subtle changes in the cortical transcriptome and epigenome. Schizophr Res 140, 175–184.

Corradini, I., Focchi, E., Rasile, M., Morini, R., Desiato, G., Tomasoni, R., Lizier, M., Ghirardini, E., Fesce, R., Morone, D., Barajon, I., Antonucci, F., Pozzi, D., Matteoli, M., 2018. Maternal Immune Activation Delays Excitatory-to-Inhibitory Gamma-Aminobutyric Acid Switch in Offspring. Biological psychiatry 83, 680–691.

Cotel, M.C., Lenartowicz, E.M., Natesan, S., Modo, M.M., Cooper, J.D., Williams, S.C., Kapur, S., Vernon, A.C., 2015. Microglial activation in the rat brain following chronic antipsychotic treatment at clinically relevant doses. Eur Neuropsychopharmacol 25, 2098–2107.

Deverman, B.E., Patterson, P.H., 2009. Cytokines and CNS development. Neuron 64, 61–78.

Elmer, B.M., Estes, M.L., Barrow, S.L., McAllister, A.K., 2013. MHCI requires MEF2 transcription factors to negatively regulate synapse density during development and in disease. J Neurosci 33, 13791–13804.

Estes, M.L., McAllister, A.K., 2014. Alterations in immune cells and mediators in the brain: it’s not always neuroinflammation! Brain Pathol 24, 623–630.

Estes, M.L., McAllister, A.K., 2015. Immune mediators in the brain and peripheral tissues in autism spectrum disorder. Nature reviews. Neuroscience 16, 469–486.

Estes, M.L., McAllister, A.K., 2016. Maternal immune activation: Implications for neuropsychiatric disorders. Science 353, 772–777.

Fatemi, S.H., Earle, J., Kanodia, R., Kist, D., Emamian, E.S., Patterson, P.H., Shi, L., Sidwell, R., 2002. Prenatal viral infection leads to pyramidal cell atrophy and macrocephaly in adulthood: implications for genesis of autism and schizophrenia. Cell Mol Neurobiol 22, 25–33.

Fillman, S.G., Cloonan, N., Catts, V.S., Miller, L.C., Wong, J., McCrossin, T., Cairns, M., Weickert, C.S., 2013. Increased inflammatory markers identified in the dorsolateral prefrontal cortex of individuals with schizophrenia. Mol Psychiatry 18, 206–214.

Garay, P.A., Hsiao, E.Y., Patterson, P.H., McAllister, A.K., 2013. Maternal immune activation causes age- and region-specific changes in brain cytokines in offspring throughout development. Brain Behav Immun 31, 54–68.

Garay, P.A., McAllister, A.K., 2010. Novel roles for immune molecules in neural development: implications for neurodevelopmental disorders. Front Synaptic Neurosci 2, 136.

Garbett, K.A., Hsiao, E.Y., Kalman, S., Patterson, P.H., Mirnics, K., 2012. Effects of maternal immune activation on gene expression patterns in the fetal brain. Translational psychiatry 2, e98.

Giovanoli, S., Engler, H., Engler, A., Richetto, J., Feldon, J., Riva, M.A., Schedlowski, M., Meyer, U., 2016. Preventive effects of minocycline in a neurodevelopmental two-hit model with relevance to schizophrenia. Translational psychiatry 6, e772.

Giovanoli, S., Notter, T., Richetto, J., Labouesse, M.A., Vuillermot, S., Riva, M.A., Meyer, U., 2015. Late prenatal immune activation causes hippocampal deficits in the absence of persistent inflammation across aging. Journal of neuroinflammation 12, 221.

Glynn, M.W., Elmer, B.M., Garay, P.A., Liu, X.B., Needleman, L.A., El-Sabeawy, F., McAllister, A.K., 2011. MHCI negatively regulates synapse density during the establishment of cortical connections. Nature neuroscience 14, 442–451.

Gore, F.M., Bloem, P.J., Patton, G.C., Ferguson, J., Joseph, V., Coffey, C., Sawyer, S.M., Mathers, C.D., 2011. Global burden of disease in young people aged 10-24 years: a systematic analysis. Lancet 377, 2093–2102.

Holmes, S.E., Hinz, R., Drake, R.J., Gregory, C.J., Conen, S., Matthews, J.C., Anton-Rodriguez, J.M., Gerhard, A., Talbot, P.S., 2016. In vivo imaging of brain microglial activity in antipsychotic-free and medicated schizophrenia: a [(11)C](R)-PK11195 positron emission tomography study. Mol Psychiatry 21, 1672–1679.

Hsiao, E.Y., Patterson, P.H., 2011. Activation of the maternal immune system induces endocrine changes in the placenta via IL-6. Brain Behav Immun 25, 604–615.

Insel, T.R., 2012. Next-generation treatments for mental disorders. Sci Transl Med 4, 155ps119.

Johnson, R.A., Wichern, D.W., 2007. Applied multivariate statistical analysis. Pearson Prentice Hall, Upper Saddle River, N.J.

Kim, J.K., Choi, B.H., Park, H.C., Park, S.R., Kim, Y.S., Yoon, S.H., Park, H.S., Kim, E.Y., Ha, Y., 2004. Effects of GM-CSF on the neural progenitor cells. Neuroreport 15, 2161–2165.

Kim, N.K., Choi, B.H., Huang, X., Snyder, B.J., Bukhari, S., Kong, T.H., Park, H., Park, H.C., Park, S.R., Ha, Y., 2009. Granulocyte-macrophage colony-stimulating factor promotes survival of dopaminergic neurons in the 1-methyl-4-phenyl-1,2,3,6-tetrahydropyridine-induced murine Parkinson’s disease model. Eur J Neurosci 29, 891–900.

Knuesel, I., Chicha, L., Britschgi, M., Schobel, S.A., Bodmer, M., Hellings, J.A., Toovey, S., Prinssen, E.P., 2014. Maternal immune activation and abnormal brain development across CNS disorders. Nature reviews. Neurology 10, 643–660.

Krieger, M., Both, M., Kranig, S.A., Pitzer, C., Klugmann, M., Vogt, G., Draguhn, A., Schneider, A., 2012. The hematopoietic cytokine granulocyte-macrophage colony stimulating factor is important for cognitive functions. Sci Rep 2, 697.

Lee, S.C., Liu, W., Brosnan, C.F., Dickson, D.W., 1994. GM-CSF promotes proliferation of human fetal and adult microglia in primary cultures. Glia 12, 309–318.

Li, Q., Cheung, C., Wei, R., Hui, E.S., Feldon, J., Meyer, U., Chung, S., Chua, S.E., Sham, P.C., Wu, E.X., McAlonan, G.M., 2009a. Prenatal immune challenge is an environmental risk factor for brain and behavior change relevant to schizophrenia: evidence from MRI in a mouse model. PLoS One 4, e6354.

Li, X., Chauhan, A., Sheikh, A.M., Patil, S., Chauhan, V., Li, X.M., Ji, L., Brown, T., Malik, M., 2009b. Elevated immune response in the brain of autistic patients. J Neuroimmunol 207, 111–116.

Livak, K.J., Schmittgen, T.D., 2001. Analysis of relative gene expression data using real-time quantitative PCR and the 2(-Delta Delta C(T)) Method. Methods 25, 402–408.

Malkova, N.V., Yu, C.Z., Hsiao, E.Y., Moore, M.J., Patterson, P.H., 2012. Maternal immune activation yields offspring displaying mouse versions of the three core symptoms of autism. Brain Behav Immun 26, 607–616.

Marin, O., 2016. Developmental timing and critical windows for the treatment of psychiatric disorders. Nat Med 22, 1229–1238.

Maung, R., Hoefer, M.M., Sanchez, A.B., Sejbuk, N.E., Medders, K.E., Desai, M.K., Catalan, I.C., Dowling, C.C., de Rozieres, C.M., Garden, G.A., Russo, R., Roberts, A.J., Williams, R., Kaul, M., 2014. CCR5 knockout prevents neuronal injury and behavioral impairment induced in a transgenic mouse model by a CXCR4-using HIV-1 glycoprotein 120. J Immunol 193, 1895–1910.

Melhem, N., Middleton, F., McFadden, K., Klei, L., Faraone, S.V., Vinogradov, S., Tiobech, J., Yano, V., Kuartei, S., Roeder, K., Byerley, W., Devlin, B., Myles-Worsley, M., 2011. Copy number variants for schizophrenia and related psychotic disorders in Oceanic Palau: risk and transmission in extended pedigrees. Biological psychiatry 70, 1115–1121.

Meyer, U., 2014. Prenatal poly(i:C) exposure and other developmental immune activation models in rodent systems. Biological psychiatry 75, 307–315.

Meyer, U., Feldon, J., Schedlowski, M., Yee, B.K., 2006a. Immunological stress at the maternal-foetal interface: a link between neurodevelopment and adult psychopathology. Brain Behav Immun 20, 378–388.

Meyer, U., Murray, P.J., Urwyler, A., Yee, B.K., Schedlowski, M., Feldon, J., 2008. Adult behavioral and pharmacological dysfunctions following disruption of the fetal brain balance between pro-inflammatory and IL-10-mediated anti-inflammatory signaling. Mol Psychiatry 13, 208–221.

Meyer, U., Nyffeler, M., Engler, A., Urwyler, A., Schedlowski, M., Knuesel, I., Yee, B.K., Feldon, J., 2006b. The time of prenatal immune challenge determines the specificity of inflammation-mediated brain and behavioral pathology. J Neurosci 26, 4752–4762.

Millan, M.J., Andrieux, A., Bartzokis, G., Cadenhead, K., Dazzan, P., Fusar-Poli, P., Gallinat, J., Giedd, J., Grayson, D.R., Heinrichs, M., Kahn, R., Krebs, M.O., Leboyer, M., Lewis, D., Marin, O., Marin, P., Meyer-Lindenberg, A., McGorry, P., McGuire, P., Owen, M.J., Patterson, P., Sawa, A., Spedding, M., Uhlhaas, P., Vaccarino, F., Wahlestedt, C., Weinberger, D., 2016. Altering the course of schizophrenia: progress and perspectives. Nat Rev Drug Discov 15, 485–515.

Mizuno, T., Zhang, G., Takeuchi, H., Kawanokuchi, J., Wang, J., Sonobe, Y., Jin, S., Takada, N., Komatsu, Y., Suzumura, A., 2008. Interferon-gamma directly induces neurotoxicity through a neuron specific, calcium-permeable complex of IFN-gamma receptor and AMPA GluR1 receptor. FASEB J 22, 1797–1806.

Mortensen, P.B., Norgaard-Pedersen, B., Waltoft, B.L., Sorensen, T.L., Hougaard, D., Torrey, E.F., Yolken, R.H., 2007. Toxoplasma gondii as a risk factor for early-onset schizophrenia: analysis of filter paper blood samples obtained at birth. Biological psychiatry 61, 688–693.

Neumann, H., Schmidt, H., Cavalie, A., Jenne, D., Wekerle, H., 1997. Major histocompatibility complex (MHC) class I gene expression in single neurons of the central nervous system: differential regulation by interferon (IFN)-gamma and tumor necrosis factor (TNF)-alpha. J Exp Med 185, 305–316.

Ozawa, K., Hashimoto, K., Kishimoto, T., Shimizu, E., Ishikura, H., Iyo, M., 2006. Immune activation during pregnancy in mice leads to dopaminergic hyperfunction and cognitive impairment in the offspring: a neurodevelopmental animal model of schizophrenia. Biological psychiatry 59, 546–554.

Paolicelli, R.C., Bolasco, G., Pagani, F., Maggi, L., Scianni, M., Panzanelli, P., Giustetto, M., Ferreira, T.A., Guiducci, E., Dumas, L., Ragozzino, D., Gross, C.T., 2011. Synaptic pruning by microglia is necessary for normal brain development. Science 333, 1456–1458.

Patterson, P.H., 2002. Maternal infection: window on neuroimmune interactions in fetal brain development and mental illness. Curr Opin Neurobiol 12, 115–118.

Pavlowsky, A., Gianfelice, A., Pallotto, M., Zanchi, A., Vara, H., Khelfaoui, M., Valnegri, P., Rezai, X., Bassani, S., Brambilla, D., Kumpost, J., Blahos, J., Roux, M.J., Humeau, Y., Chelly, J., Passafaro, M., Giustetto, M., Billuart, P., Sala, C., 2010. A postsynaptic signaling pathway that may account for the cognitive defect due to IL1RAPL1 mutation. Curr Biol 20, 103–115.

Piontkewitz, Y., Assaf, Y., Weiner, I., 2009. Clozapine administration in adolescence prevents postpubertal emergence of brain structural pathology in an animal model of schizophrenia. Biological psychiatry 66, 1038–1046.

Prinz, M., Priller, J., 2014. Microglia and brain macrophages in the molecular age: from origin to neuropsychiatric disease. Nature reviews. Neuroscience 15, 300–312.

Re, F., Belyanskaya, S.L., Riese, R.J., Cipriani, B., Fischer, F.R., Granucci, F., Ricciardi-Castagnoli, P., Brosnan, C., Stern, L.J., Strominger, J.L., Santambrogio, L., 2002. Granulocyte-macrophage colony-stimulating factor induces an expression program in neonatal microglia that primes them for antigen presentation. J Immunol 169, 2264–2273.

Reisinger, S.N., Kong, E., Khan, D., Schulz, S., Ronovsky, M., Berger, S., Horvath, O., Cabatic, M., Berger, A., Pollak, D.D., 2016. Maternal immune activation epigenetically regulates hippocampal serotonin transporter levels. Neurobiol Stress 4, 34–43.

Schabitz, W.R., Kruger, C., Pitzer, C., Weber, D., Laage, R., Gassler, N., Aronowski, J., Mier, W., Kirsch, F., Dittgen, T., Bach, A., Sommer, C., Schneider, A., 2008. A neuroprotective function for the hematopoietic protein granulocyte-macrophage colony stimulating factor (GMCSF). J Cereb Blood Flow Metab 28, 29–43.

Schafer, D.P., Lehrman, E.K., Kautzman, A.G., Koyama, R., Mardinly, A.R., Yamasaki, R., Ransohoff, R.M., Greenberg, M.E., Barres, B.A., Stevens, B., 2012. Microglia sculpt postnatal neural circuits in an activity and complement-dependent manner. Neuron 74, 691–705.

Schafer, D.P., Lehrman, E.K., Stevens, B., 2013. The “quad-partite” synapse: microglia-synapse interactions in the developing and mature CNS. Glia 61, 24–36.

Shi, L., Smith, S.E., Malkova, N., Tse, D., Su, Y., Patterson, P.H., 2009. Activation of the maternal immune system alters cerebellar development in the offspring. Brain Behav Immun 23, 116–123.

Shin Yim, Y., Park, A., Berrios, J., Lafourcade, M., Pascual, L.M., Soares, N., Yeon Kim, J., Kim, S., Kim, H., Waisman, A., Littman, D.R., Wickersham, I.R., Harnett, M.T., Huh, J.R., Choi, G.B., 2017. Reversing behavioural abnormalities in mice exposed to maternal inflammation. Nature 549, 482–487.

Smith, S.E., Li, J., Garbett, K., Mirnics, K., Patterson, P.H., 2007. Maternal immune activation alters fetal brain development through interleukin-6. J Neurosci 27, 10695–10702.

Song, J.H., Wang, C.X., Song, D.K., Wang, P., Shuaib, A., Hao, C., 2005. Interferon gamma induces neurite outgrowth by up-regulation of p35 neuron-specific cyclin-dependent kinase 5 activator via activation of ERK1/2 pathway. J Biol Chem 280, 12896–12901.

Sorensen, H.J., Mortensen, E.L., Reinisch, J.M., Mednick, S.A., 2009. Association between prenatal exposure to bacterial infection and risk of schizophrenia. Schizophr Bull 35, 631–637.

Trepanier, M.O., Hopperton, K.E., Mizrahi, R., Mechawar, N., Bazinet, R.P., 2016. Postmortem evidence of cerebral inflammation in schizophrenia: a systematic review. Mol Psychiatry 21, 1009–1026.

Valnegri, P., Montrasio, C., Brambilla, D., Ko, J., Passafaro, M., Sala, C., 2011. The X-linked intellectual disability protein IL1RAPL1 regulates excitatory synapse formation by binding PTPdelta and RhoGAP2. Hum Mol Genet 20, 4797–4809.

Vargas, D.L., Nascimbene, C., Krishnan, C., Zimmerman, A.W., Pardo, C.A., 2005. Neuroglial activation and neuroinflammation in the brain of patients with autism. Ann Neurol 57, 67–81.

Victorio, S.C., Cartarozzi, L.P., Hell, R.C., Oliveira, A.L., 2012. Decreased MHC I expression in IFN gamma mutant mice alters synaptic elimination in the spinal cord after peripheral injury. Journal of neuroinflammation 9, 88.

Vuillermot, S., Weber, L., Feldon, J., Meyer, U., 2010. A longitudinal examination of the neurodevelopmental impact of prenatal immune activation in mice reveals primary defects in dopaminergic development relevant to schizophrenia. J Neurosci 30, 1270–1287.

Wei, H., Zou, H., Sheikh, A.M., Malik, M., Dobkin, C., Brown, W.T., Li, X., 2011. IL-6 is increased in the cerebellum of autistic brain and alters neural cell adhesion, migration and synaptic formation. Journal of neuroinflammation 8, 52.

Winter, C., Reutiman, T.J., Folsom, T.D., Sohr, R., Wolf, R.J., Juckel, G., Fatemi, S.H., 2008. Dopamine and serotonin levels following prenatal viral infection in mouse--implications for psychiatric disorders such as schizophrenia and autism. Eur Neuropsychopharmacol 18, 712–716.

Yirmiya, R., Goshen, I., 2011. Immune modulation of learning, memory, neural plasticity and neurogenesis. Brain Behav Immun 25, 181–213.

Zhan, Y., Paolicelli, R.C., Sforazzini, F., Weinhard, L., Bolasco, G., Pagani, F., Vyssotski, A.L., Bifone, A., Gozzi, A., Ragozzino, D., Gross, C.T., 2014. Deficient neuron-microglia signaling results in impaired functional brain connectivity and social behavior. Nature neuroscience 17, 400–406.

Zhou, M., Greenhill, S., Huang, S., Silva, T.K., Sano, Y., Wu, S., Cai, Y., Nagaoka, Y., Sehgal, M., Cai, D.J., Lee, Y.S., Fox, K., Silva, A.J., 2016. CCR5 is a suppressor for cortical plasticity and hippocampal learning and memory. Elife 5.

